# Gain-of-function study reveals the pleiotropic roles of serine protease HtrA in *Borrelia burgdorferi*

**DOI:** 10.1101/2024.08.28.610130

**Authors:** Kai Zhang, Ching Wooen Sze, Hang Zhao, Jun Liu, Chunhao Li

## Abstract

High-temperature requirement protease A (HtrA) is a family of serine proteases degrading misfolded and damaged proteins that are toxic to bacteria. The Lyme disease agent *Borrelia burgdorferi* encodes a single HtrA (BbHtrA). Previous studies have shown that BbHtrA is a key virulence determinant of *B. burgdorferi* as a deletion mutant of *htrA* (*ΔhtrA*) fails to establish infection in mice. However, previous complementation could only restore protein expression but not infectivity in mice. In this report, we first identify the native promoter of BbHtrA which allows us to construct a fully complemented *ΔhtrA* strain. Follow up promoter activity analysis reveals that BbHtrA is likely dually regulated by the house keeping sigma factor RpoD and the alternative sigma factor RpoS. The *ΔhtrA* mutant exhibits growth defect upon entering the mid-log to stationary phase especially at high temperatures. Microscopic analysis further demonstrates that the absence of *htrA* induces extensive cell death. Additionally, the *ΔhtrA* mutant has defects in cell locomotion as the expression of several key chemotaxis proteins are significantly downregulated. Cryo-electron tomography imaging of *htrA* mutant further reveals that deletion of *htrA* disrupts flagellar homeostasis. The failure of *ΔhtrA* to establish an infection in mice is likely due to repressed expression of BosR and RpoS at the transcriptional level which ultimately causes dysregulation of the RpoS-induced virulence factors. Collectively, we conclude that the expression of *htrA* is finely tuned which is critical for its pleiotropic roles in the regulation of motility, stress response, and virulence gene expression in *B. burgdorferi*.

**IMPORTANCE:** Lyme borreliosis is the most commonly reported vector-borne illnesses in the United States, which is caused by *Borrelia burgdorferi.* As the enzootic pathogen alternates between the tick vector and mammalian hosts, adaptation to drastically different growth milieu is imperative to its survival. Hence, robust alteration of gene expression and proper quality control on protein synthesis and turnover are pivotal for its fitness. The family of HtrA serine proteases is mainly responsible for the maintenance of protein homeostasis particularly under stressful conditions. The significance of this report is to decode how BbHtrA contributes to the fitness of *B. burgdorferi*. BbHtrA is essential for mammalian host infection but little is known about its regulatory mechanism as well as its contribution to the virulence of *B. burgdorferi*. By deciphering the regulatory elements involved in the expression of BbHtrA, we are one step closer to comprehending its significance in the pathophysiology of *B. burgdorferi*.

## INTRODUCTION

Lyme disease (LD) is a vector-borne infectious disease caused by the bacterium *Borrelia burgdorferi* (recently reassigned to the genus *Borreliella*) and transmitted to humans through the bite of infected *Ixodes scapularis* ticks (1–4). In the United States, it estimates that there are 476,000 new cases annually, making LD one of the most common tick-borne illnesses in the country (5–7). Early-stage LD can be easily treated with antibiotics, however, unrecognized late stage LD can have debilitating long-term health effects, such as chronic arthritis and cognitive impairment, which can manifest after months or even years of latent infection (8, 9). In addition, some patients who are non-responsive to antibiotic therapy can develop post-treatment Lyme disease syndrome (PTLDS) which poses a significant challenge not only to the individual but also presents as a big economic burden on the U.S. healthcare system (10).

As *B. burgdorferi* alternates between the arthropod vector and mammalian host during its complex enzootic cycle, robust transcriptomic, proteomic, and metabolic alterations are vital for its adaptation and survival under distinct host environments (2, 11–14). For instance, many antigens are differentially expressed under tick- and mammalian-milieu to assist in invasion, dissemination, immune evasion, and metabolic adaptation (1, 11–13, 15, 16). When bacteria are exposed to drastic changes in the environment, such as during the transition of *B. burgdorferi* between ticks and mammals, protein homeostasis needs to be maintained through several coordinated processes such as protein synthesis, degradation, and chaperone-mediated quality control to ensure proper protein folding and functioning under stressful conditions (17, 18).

Proteases are enzymes that play significant roles in maintaining protein homeostasis in bacteria through the removal of misfolded and damaged proteins via regulated protein degradation (19, 20). These processes are important for bacterial protein quality control by specifically targeting selected proteins for degradation to prevent toxic accumulation and cellular stresses. One important family of such proteases is the HtrA (<u>H</u>igh <u>T</u>emperature <u>R</u>equirement <u>A</u>) serine protease (21–23). HtrA belongs to a highly conserved family of serine proteases that are found in higher life forms such as humans as well as prokaryotes like bacteria (24–26). In *Escherichia coli*, three homologs of HtrA proteins, DegP, DegQ, and DegS, have been well studied and served as the prototype for understanding the function of HtrA enzymes (27–31). The coordinated action of these three HtrA proteins in *E. coli* prevents stress-induced aggregation of misfolded proteins (26).

A HtrA homolog has been identified in *B. burgdorferi* (BbHtrA) and its substrate specificity and contribution to the pathogenesis of *B. burgdorferi* have been characterized by several reports (32–40). BbHtrA has been shown to degrade host extracellular matrix proteins and fibronectin, which may contribute to the dissemination process of the spirochetes during infection (34, 35). Additionally, several proteins essential for *B. burgdorferi* chemotaxis/motility and infectivity in mammals have also been identified as the substrates of BbHtrA (32, 36, 38, 40). *In vivo* mouse infection study revealed that BbHtrA is absolutely required for mammalian infectivity presumably to assist in the proteolytic processing of BB0323, a virulence determinant of *B. burgdorferi* (41). However, the pathogenic defect of BbHtrA mutant in mice was not confirmed through gain-of-function study due to technical constraints (38). In addition, how BbHtrA expression is regulated and the underlying mechanism contributing to *B. burgdorferi* infectivity has yet to be determined. In this report, we sought to answer these questions through the complementation of our BbHtrA mutant by first identifying the endogenous promoter of this gene. The impact of BbHtrA on *B. burgdorferi* mouse infection was then re-evaluated using our complemented strains and the underlying regulatory mechanism of BbHtrA on *B. burgdorferi* pathogenesis was further investigated in detail.

## RESULTS

### Isolation of the *htrA* mutant and its isogenic complemented strains

A *htrA* (*bb0104*) in frame deletion mutant, *ΔhtrA,* was constructed in B31 A3-68, a low-passage virulent *B. burgdorferi* strain as described in our previous report (40). Initial complementation attempts were performed *in trans* with pBSV2G (42) and pBBE22G (43) shuttle vectors using two well-characterized *B. burgdorferi* promoters, *pflgB* (44) and *pflaB* (45), generating three complemented vectors: pflgBbb0104/pBSV2G, pflaBbb0104/pBSV2G, and pflgBbb0104/pBBE22G (**Fig. 1A**). With these vectors, three different complemented strains (*ΔhtrA^C1^*, *ΔhtrA^C2^,* and *ΔhtrA^C3^*) were generated and then tested in mice for infectivity. Surprisingly, none of these complemented strains was able to establish infection in mice (Table 1) even though the expression of BbHtrA was successfully restored (**Fig. 1B**) and that all plasmids essential for infectivity are retained (**Fig. 1D**). A similar phenotype was previously reported by Ye *et al.* (38).

**Figure 1.**
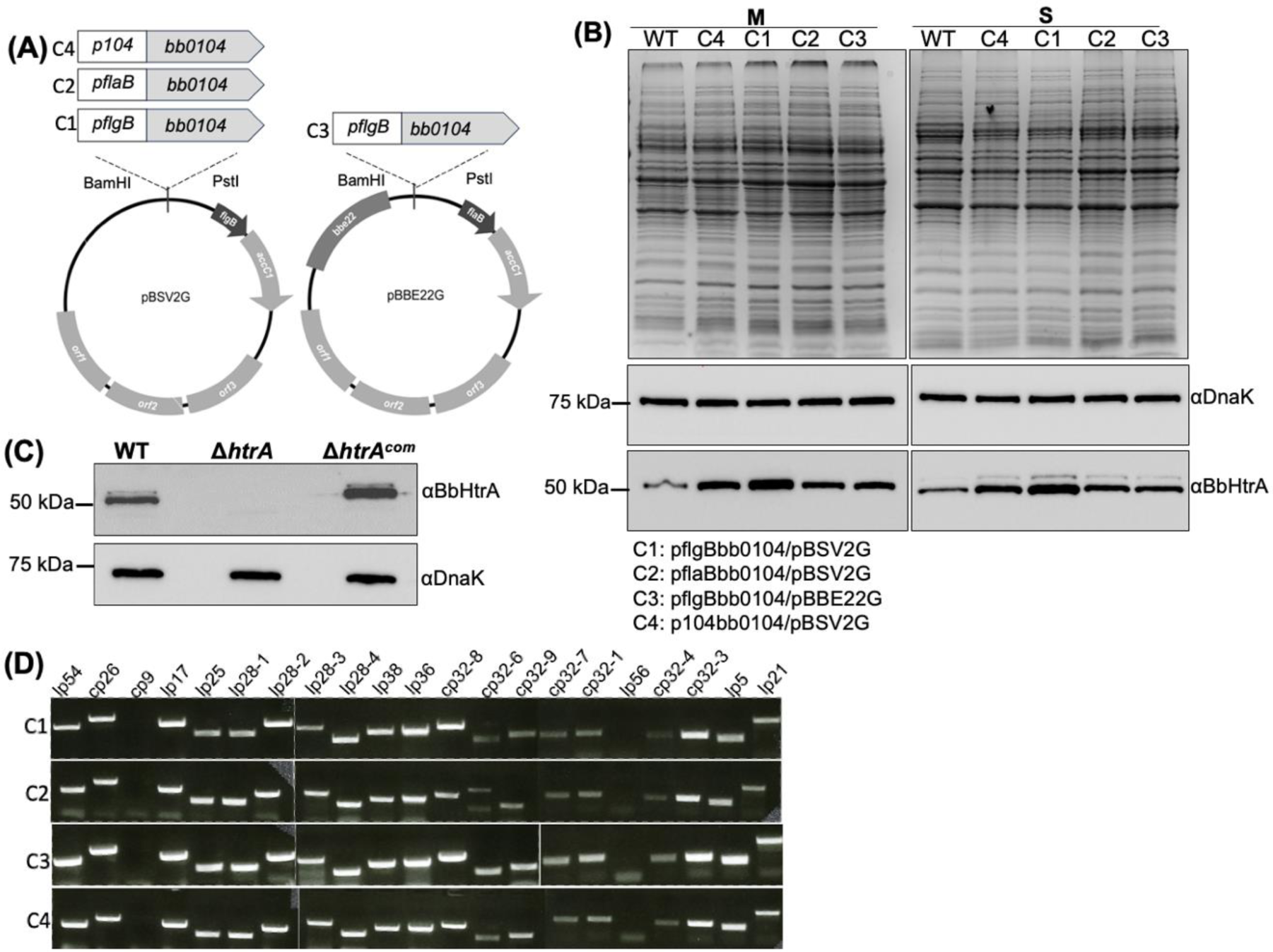
Isolation and characterization of BbHtrA complemented strains. **(A)** Construction of *bb0104* trans-complementation plasmids. Three different promoters including *flgB* (*pflgB*), *flaB* (*pflaB*) and the native promoter upstream *bb0104* (*p104*) were fused to *bb0104* and then cloned into the shuttle vector pBSV2G or pBBE22G, yielding four vectors (pflgBbb0104/pBSV2G, pflaBbb0104/pBSV2G, pflgBbb0104/pBBE22G, and p104bb0104/pBSV2G) for the trans-complementation of *ΔhtrA*. The resulted corresponding complemented strains were designated as C1, C2, C3, and C4, respectively. **(B)** Immunoblotting analysis of WT and four BbHtrA complemented strains harvested at mid-log (M) and stationary (S) growth phase. For this experiment, the same amounts of WT and complemented whole cell lysates were analyzed by SDS-PAGE and then probed with BbHtrA and DnaK antibodies. DnaK was used as loading control. **(C)** Immunoblotting analysis of *ΔhtrA* and a fully complemented strain *ΔhtrA^com^*. Same amounts of WT, *ΔhtrA*, and fully complemented *ΔhtrA^com^*whole cell lysates were analyzed and probed with antibodies against BbHtrA and DnaK. DnaK was used as a loading control. **(D)** Detecting the plasmid profiles of four *htrA* complemented strains by PCR. This study was performed as previously described (86). Of note, B31 A3-68, the parental strain was used to construct *ΔhtrA* and the four isogenic complemented strains, does not contain the plasmid lp56 (83).

**Table 1.** BbHtrA is required for infectivity in BALB/c mice.

| *B. burgdorferi* strains | No. of positive cultures/total no. of specimens examined |  |  |  |  |
| --- | --- | --- | --- | --- | --- |
|  | Ear | Skin | Joint | Heart | Spleen |
| B31 A3-68 | 12/12 | 12/12 | 12/12 | 12/12 | 12/12 |
| $\Delta htrA$ | 0/12 | 0/12 | 0/12 | 0/12 | 0/12 |
| $\Delta htrA^{C1}$ | 0/3 | 0/3 | 0/3 | 0/3 | 0/3 |
| $\Delta htrA^{C2}$ | 0/3 | 0/3 | 0/3 | 0/3 | 0/3 |
| $\Delta htrA^{C3}$ | 0/3 | 0/3 | 0/3 | 0/3 | 0/3 |
| $\Delta htrA^{com}$ | 3/3 | 3/3 | 3/3 | 3/3 | 3/3 |

C1: pflgBbb0104/pBSV2G; C2: pflaBbb0104/pBSV2G; C3: pflgBbb0104/pBBE22G

### Mapping *bb0104* promoter

The unsuccessful restoration of infectivity in three complement strains above suggests that the expression timing and level of BbHtrA might be critical for the pathogenicity of *B. burgdorferi* which needs to be tightly regulated *in vivo*. As both *flaB* and *flgB* promoters are strong and constitutively active promoters (44, 46, 47), the level of BbHtrA may be over-induced or failed to be turned off when necessary *in vivo*. Hence identifying the native promoter for *bb0104* is necessary to fully comprehend the regulation and contribution of BbHtrA to the life cycle of *B. burgdorferi*. To precisely map the transcriptional start site (TSS) of *htrA*, 5′-RACE was performed using primer targeted to the open reading frame (*orf*) of *bb0104* (Table 2). The obtained 5’-RACE amplicon was subjected to DNA sequencing and the result showed that the amplicon generated by the primer started at a “G” nucleotide which is located within *bb0105*, 122 bp from its stop codon and 191 bp from the start codon of *htrA* (**Fig. 2A**). This result is also in congruency with the *B. burgdorferi* 5′ end transcriptome Interactive Genomics Research database (48). We searched the upstream region of TSS for promoter consensus sequences that have been characterized in *B. burgdorferi*, such as BosR, RpoS, RpoD, and RpoN but did not find any. However, transcriptional reporter assays using *lacZ* in *E. coli* confirmed the presence of a functional promoter (*p104*, promoter of *bb0104*) within the 300 bp sequence upstream of *htrA*. As shown in Fig 2C, the average *β*-galactosidase activity for *p104* is 5559.89 ± 1275.05 Miller Units. No signal was detected from cells carrying only the empty vector *pRS414*, while a strong signal was detected from our positive control *flaB* promoter, *pflaB* (13558.54 ± 1407.30 Miller units) (40). Collectively, 5’-RACE and transcriptional reporter assays indicate that the upstream region of *htrA* contains an active promoter.

**Figure 2.**
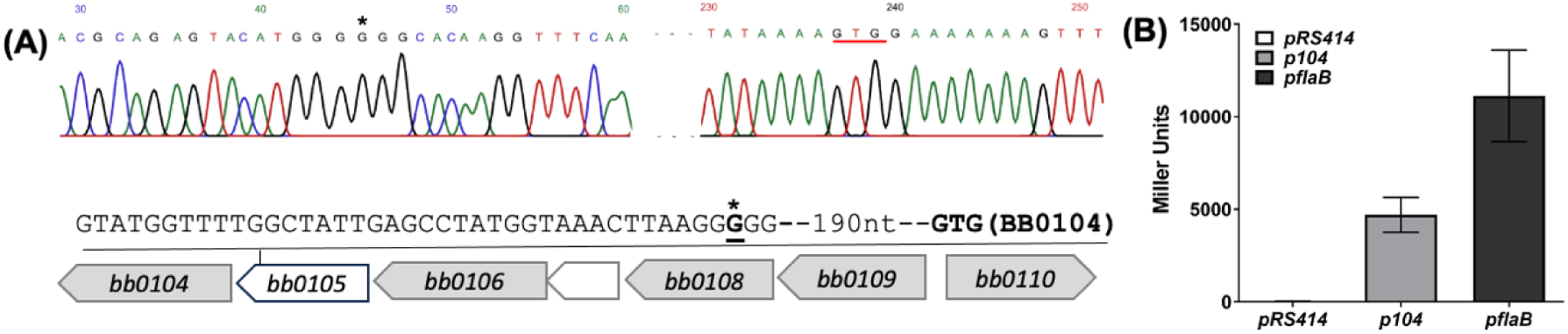
Mapping *bb0104* promoter. **(A)** 5′-RACE analysis was performed using primer targeted to *bb0104* to identify the transcriptional start site (TSS). The TSS of *bb0104* was mapped to a G nucleotide 192 base pair (bp) from the start codon of *bb0104* located inside the open reading frame *(orf)* of *bb0105.* A representative chromatogram of the sequencing result is shown. Asterisk marks the TSS, red underlined GTG is the start codon of *bb0104*. **(B)** *β*-galactosidase assay using *bb0104* promoter (*p104*). For this assay, the upstream region (300 bp) of *bb0104* was cloned into pRS414 vector. The empty vector pRS414, the vector with *p104*, and a positive control pRS414 with *flaB* promoter (*pflaB*) were electroporated into *E. coli* and the promoter activity was measured by quantifying the *β*-galactosidase activity. The results were expressed as the average Miller units of triplicate samples from three independent experiments ± standard errors of the means (SEM).

**Table 2.**
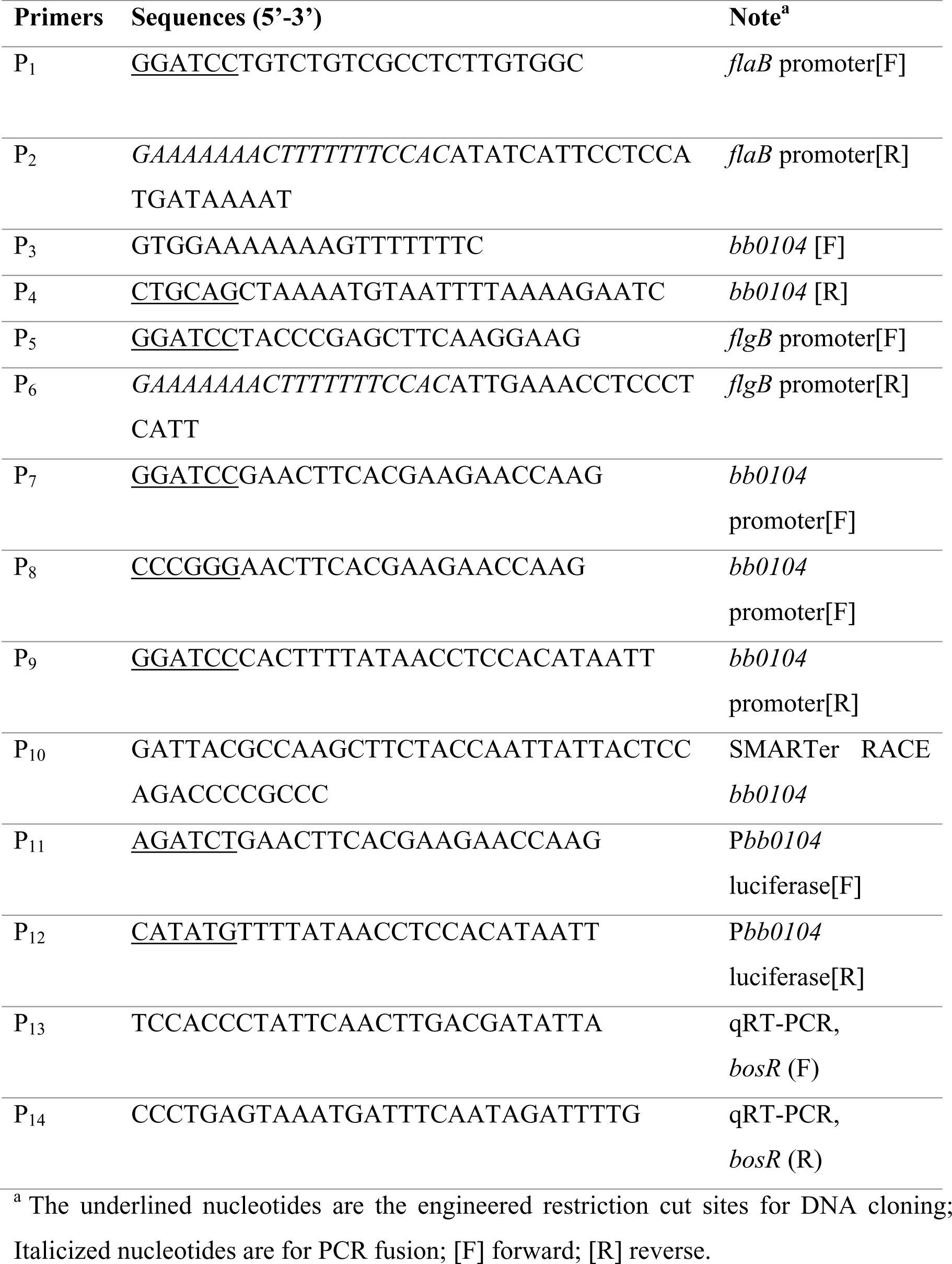
Oligonucleotide primers used in this study.

### Reduced *htrA* promoter activities in the absence of BosR and RpoS

Based on our 5′-RACE result, we were not able to locate a well-conserved -10 Pribnow box and -35 sequence. To investigate how BbHtrA expression is regulated, the identified *htrA* promoter (*p104*) was ligated to a luciferase reporter vector, pJSB161 (49) and transformed into WT, *bosR* (*ΔbosR*), and *rpoS* (*ΔrpoS*) mutants as described previously (50). Cells were cultivated at 37°C/pH 6.8 for up to 12 days to mimic the mammalian host environment. Cell counts and luciferase reporter activity was quantified every two days until the cells entered the stationary phase. The luciferase readings were normalized against WT on day 2 since there was no detectable signals from cells carrying the empty vector pJSB161. In the absence of BosR and RpoS, the level of luciferase expression driven by *p104* was significantly reduced when compared to the WT cells carrying the same plasmid. A gradual increase in the luciferase activity could be seen over time in the WT cells with the peak expression occurring at the late-log to stationary phase. The same pattern was seen in both *ΔbosR* and *ΔrpoS* mutants but at a significantly lower level than in the WT (**Fig. 3A**). This result implies that the optimal activity of *p104* requires the presence of RpoS and/or BosR. In congruence with this, qRT-PCR analysis revealed significantly lower *htrA* transcripts in *ΔbosR* and *ΔrpoS* as compared to the WT at the mid-log growth phase which was partially restored upon entering the stationary phase (**Fig. 3B**).

**Figure 3.**
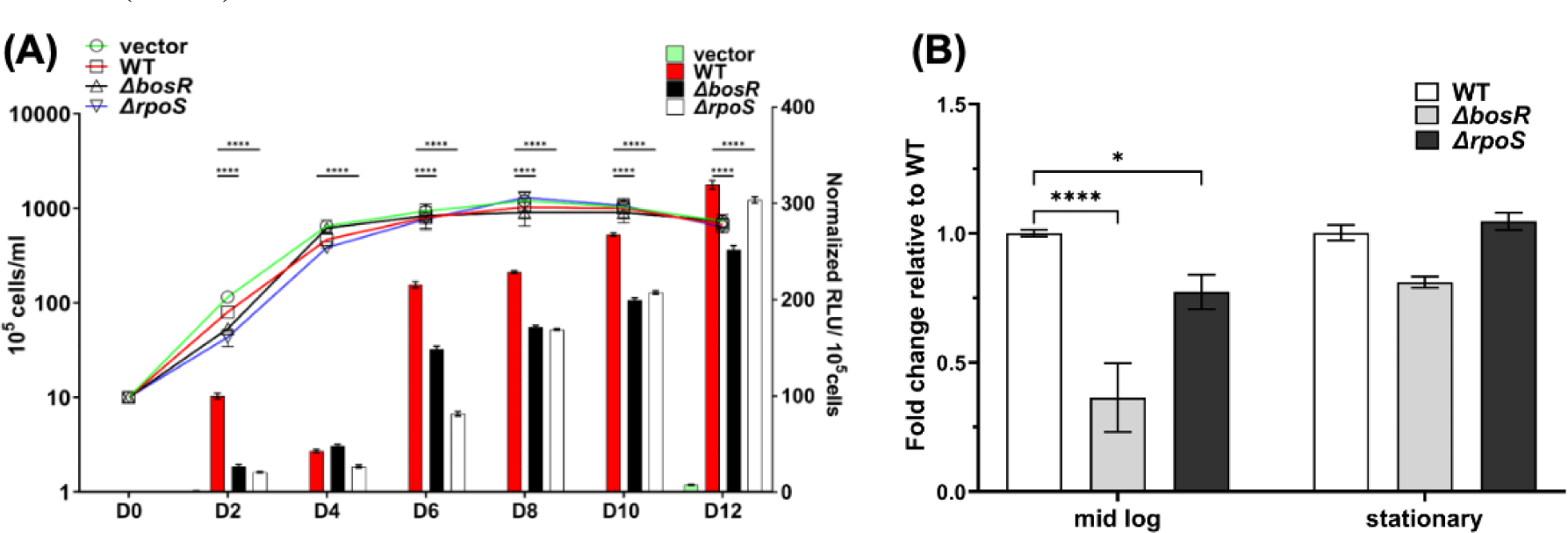
Dual regulation of *htrA*. **(A)** The *htrA* promoter activity is significantly reduced in the absence of BosR, and RpoS. For this experiment, the native promoter of *bb0104* (*p104*) identified through 5′-RACE was cloned into pJSB161, a modified luciferase reporter construct (49). The resulted constructs were transformed into *B. burgdorferi* WT, *ΔbosR*, and *ΔrpoS* strains, respectively. Cell growth at 37°C 5% CO_2_ was monitored using Petroff Hausser counting chamber and luciferase activity was quantified using Varioskan LUX multimode microplate reader over 12 days. Samples were collected for luciferase activity reading every 2 days. Data was expressed as relative light units per 10^5^ cells (RLU/10^5^ cells). Statistical analysis was performed between WT, *ΔbosR*, and *ΔrpoS* using two-way ANOVA. **(B)** Reduced *htrA* transcript in *ΔbosR* and *ΔrpoS*. The transcript level of *htrA* in *ΔbosR* and *ΔrpoS* mutants was compared to the WT at mid-log and stationary growth phase. The mean of two biological replicates is shown here. The significance of the differences between experimental groups was evaluated using two-way ANOVA (*P* value < 0.05) asterisk (*) denotes a statistically significant difference.

### Complementation using the native promoter restores *ΔhtrA* infectivity in mice

With the successful mapping of the *htrA* native promoter (*p104*), we attempted to complement *ΔhtrA* using p104bb0104/pBSV2G (**Fig. 1A**). Complementation successfully restored the expression of BbHtrA as shown by immunoblots (**Fig. 1B, C**). One clone with the same plasmid profile as the WT was selected and annotated as *ΔhtrA^com^* (**Fig. 1D**). Using *ΔhtrA^com^*, we repeated the mouse infection study as described above and successfully isolated spirochetes from various organs of the infected mice (i.e., ear, skin, joint, heart and spleen) to the same extend as the WT-infected mice (Table 1). This result confirms that complementation of *ΔhtrA* using the native promoter that we have identified through 5′-RACE can fully restore the infectivity of the mutant. This was the first report of *htrA* mutant complementation where infectivity was fully recovered.

To understand how complementation using the native promoter was able to restore the infectivity of *ΔhtrA* but not with *flgB* or *flaB* promoters (**Fig. 1B**), the expression profile of several key virulence factors in *B. burgdorferi* were compared among the four complemented strains. Immunoblotting analysis showed that all four complemented strains have restored BbHtrA expression but at varying levels. Complemented strains *ΔhtrA^C1^* (C1) and *ΔhtrA^C2^* (C2) have the highest level of BbHtrA expressed as compared to *ΔhtrA^C3^* (C3) and *ΔhtrA^com^* (C4) at both the mid-log and stationary growth phase (**Fig. 4**). Additionally, the expression of several key virulence factors (e.g., BBK32, BosR, and DbpA) also varied among the four strains. This result implies that timely expression as well as the level of BbHtrA are crucial in the infectivity of *B. burgdorferi*. Similarly, the shuttle vectors used for complementation can also contribute to the differential expression of virulence determinants as observed in C1 vs C3. pBBE22G is derived from pBSV2G where *bbe22*, a gene encoding a nicotinamidase (PncA) was inserted into the shuttle vector pBSV2G, forming pBBE22G (43). This plasmid is often used to complement *B. burgdorferi* strains that have lost the lp25 plasmid to restore their infectivity in mammalian host. Inclusion of *bbe22* in the complementation vector appears to have a positive effect on the expression of BosR and RpoS leading to enhance expression of RpoS-regulon in C3, such as OspC and DbpA (**Fig. 4**).

**Figure 4.**
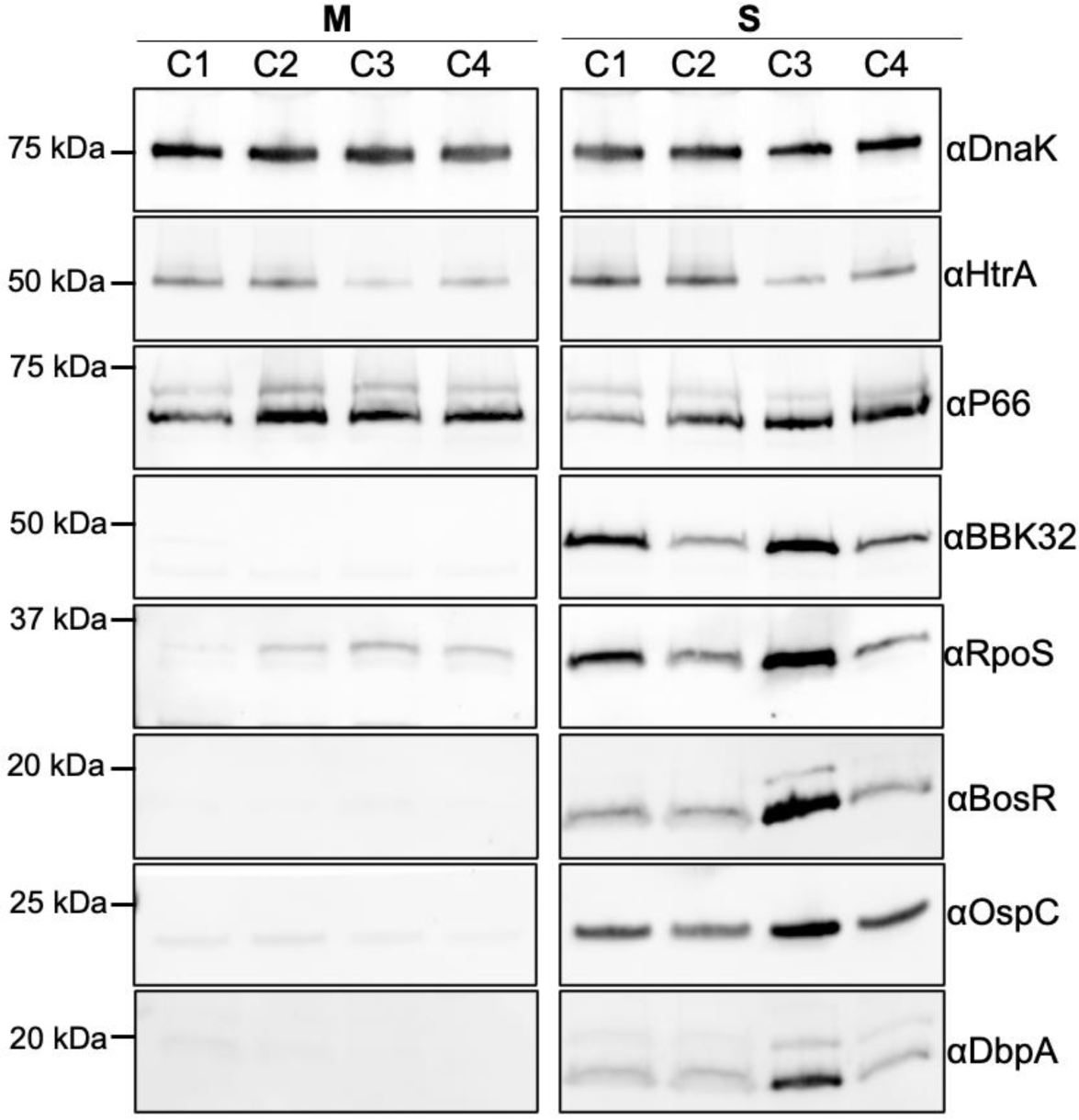
BbHtrA affects the expression of key virulence factors. Four Δ*htrA* complemented strains (C1, C2, C3, and C4) were cultivated at 37°C/pH 6.8 to mimic the mammalian host condition and harvested at mid-log (M) (∼10^6^ cells/ml) or stationary phase (S) (∼10^8^ cells/ml) for immunoblotting analysis. For this experiment, similar amounts of whole cell lysates were analyzed by stain-free SDS-PAGE and then subjected to immunoblots using specific antibodies against BbHtrA, P66 (98), BBK32 (99), RpoS, BosR, OspC, and DbpA (73). DnaK was used as an internal control, as previously described (85).

### Differential expression of BbHtrA

Luciferase reporter assay and qRT-PCR indicated that the expression of *htrA* can be regulated by both BosR and RpoS which may explain why only the native promoter of BbHtrA is able to restore the infectivity of *ΔhtrA*. To further delineate the expression pattern of BbHtrA, wild-type *B. burgdorferi* were cultivated under conditions to mimic the unfed tick (23°C/pH 7.4), mammalian host environment (37°C/pH 6.8), and laboratory culture condition (34°C/pH 7.4). The expression pattern of BbHtrA was assessed at both the mid-log and stationary growth phases. At the protein level, BbHtrA expression was the highest under 23°C, followed by 34°C and 37°C under both the mid-log and stationary growth phases (**Fig. 5A**). Relative to DnaK, the level of BbHtrA was approximately 4-fold lower at 37°C as compared to 23°C (**Fig. 5B**). This result further emphasizes that BbHtrA is differentially expressed under the regulation of its native promoter.

**Figure 5.**
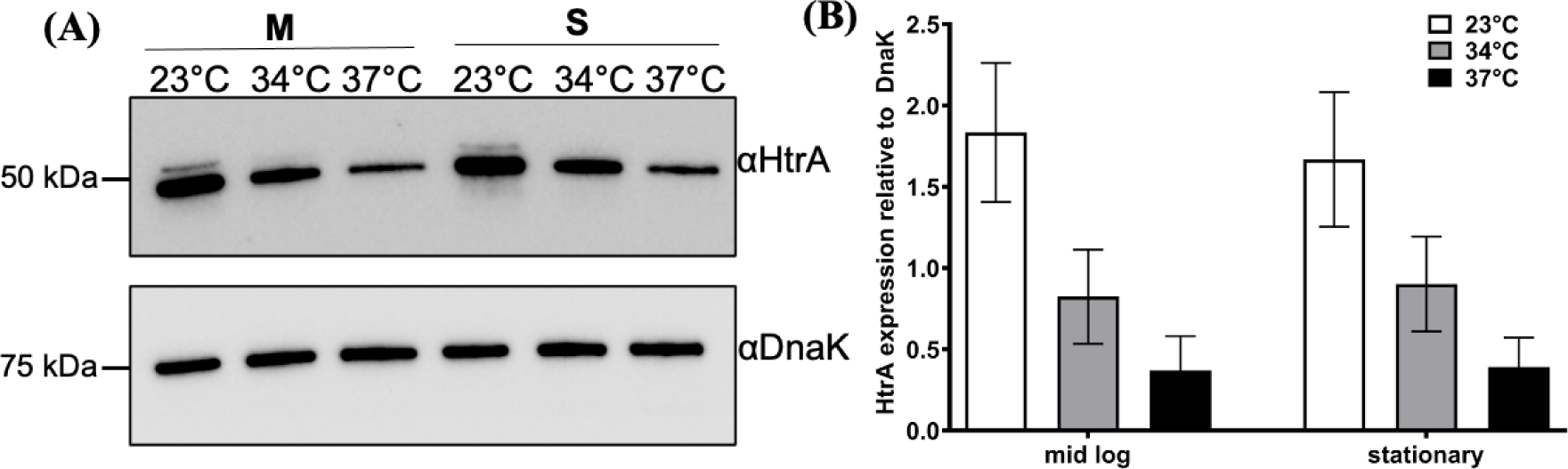
BbHtrA is differentially expressed. **(A)** Immunoblot analysis of BbHtrA expression pattern at 23°C/pH 7.4, 34°C/pH 7.4, and 37°C/pH 6.8. Wild-type *B. burgdorferi* was cultivated under various conditions and harvested for immunoblot analysis at mid-log (M) and stationary (S) growth phase. The expression level of BbHtrA was detected and DnaK was used as loading control. **(B)** Normalized expression level of BbHtrA relative to DnaK. BbHtrA signals from the immunoblot analysis were quantified using Image Lab software and normalized to the intensity of their respective DnaK level. Data from two replicates are expressed as mean expression ± SEM relative to DnaK.

### The *ΔhtrA* mutant has impaired growth upon entering the mid-log phase and at elevated temperature

Previous study revealed that deletion of *htrA* in the *B. burgdorferi* strain 297 clone AH130 background impairs its ability to grow at elevated temperature (38). Since our *htrA* mutant was constructed in B31 A3-68 strain, we repeated a similar growth analysis using WT, *ΔhtrA*, and *ΔhtrA^com^* under various conditions to mimic the unfed tick (UF, 23°C/pH7.4), routine laboratory condition (34°C/pH7.4), and mammalian phase (37°C/pH6.8). Briefly, 10^5^ cells of stationary phase seed cultures were inoculated into fresh BSK-II medium adjusted to the specified pH as indicated, and cell counts were performed every 1-3 days until stationary phase (∼10^8^ cells/ml). The *ΔhtrA* mutant grew similarly as the WT and *ΔhtrA^com^* during the early growth phase. However, upon entering the mid-log stage, the mutant started to exhibit reduced growth rates under all three conditions which persisted into the stationary phase. The observed growth defect was most prominent at 37°C where a ∼1 log difference in cell numbers was observed between the WT and *ΔhtrA*. Complementation fully reversed the growth defect to the wild-type level under all growth conditions (**Fig. 6A**), implicating that the reduced growth rate observed in *ΔhtrA* is due to the deletion of *htrA*. Significant difference in the cell numbers between *ΔhtrA* and the WT was also observed in cultures cultivated under UF and routine laboratory condition albeit to a lesser extends than under mammalian-like condition (∼ 0.6 and 0.3 log, respectively) (**Fig. 6A**).

**Figure 6.**
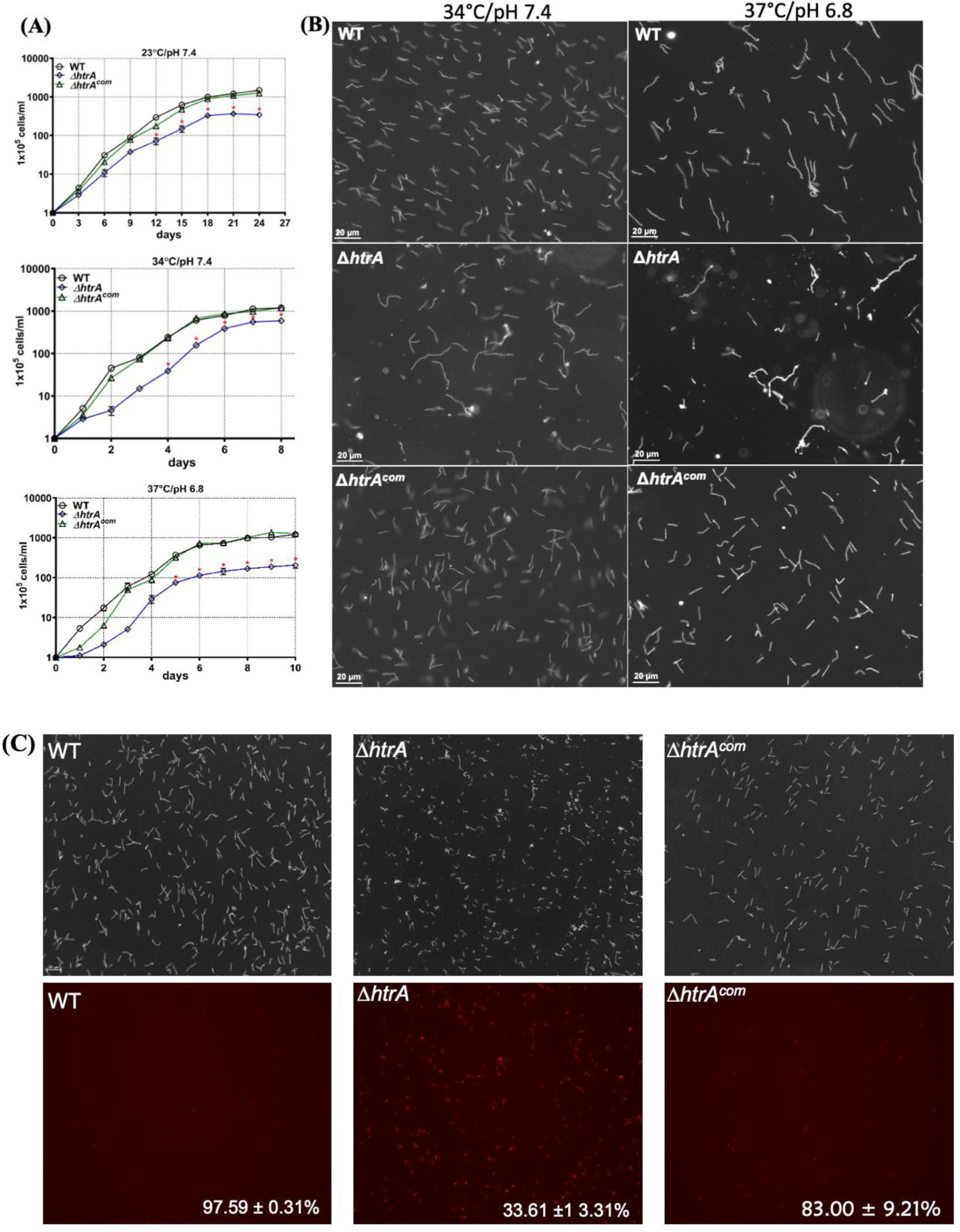
*ΔhtrA* has growth defect upon entering the mid-log to stationary growth phase with altered cell morphology. **(A)** Growth analysis of WT, *ΔhtrA*, and *ΔhtrA^com^* at 23°C/pH 7.4, 34°C/pH 7.4, and 37°C/pH 6.8. Growth curve analysis was carried out to determine if BbHtrA affects cell growth. 10^5^ cells/ml of bacteria were inoculated into 10ml BSK-II medium and cultivated under the indicated temperatures and conditions. Cell numbers were enumerated between 1-3 days until cells entered the stationary phase. Cell counting was repeated in triplicate with at least three independent samples, and the results are expressed as means ± SEM. **(B)** *ΔhtrA* has altered cell morphology and enhanced cell death at elevated temperature. Microscopic images of WT, *ΔhtrA*, and *ΔhtrA^com^* taken at the end of the growth curve analysis under 34°C/pH 7.6 and 37°C/pH 6.8. **(C)** Enhanced cell death in *ΔhtrA*. WT, *ΔhtrA*, and *ΔhtrA^com^* were cultivated at 37°C/pH 6.8. Upon entering the stationary growth phase, cells were stained with propidium iodide to identify dead cells. The percentage of dead cells was calculated from randomly chosen fields using the number of propidium iodide positive cells over total cell number under dark field. Images were taken under 200 x magnification using a Zeiss Axiostar Plus microscope. Scale bars represent 20 μm.

### Enhanced cell death observed in *ΔhtrA* at an elevated temperature

The above growth analysis indicates that BbHtrA is necessary for the growth of *B. burgdorferi* after the mid-log phase especially at an elevated temperature (**Fig. 6A**). We observed that the reduced cell number in *ΔhtrA* is due to excessive cell death (**Fig. 6B**). Live and dead staining using propidium iodide of stationary phase cells revealed an average live cell rate of 97.59 ± 0.31 % in the WT, while a drastic decrease to 33.61 ± 13.31 % was observed in *ΔhtrA.* The survival rate was restored in *ΔhtrA^com^*to 83 ± 9.21 % (**Fig. 6C**), further confirming that the deletion of *htrA* is detrimental to *B. burgdorferi* growth at elevated temperature upon entering the mid-log to stationary phase. Additionally, extensive morphological changes were observed in *ΔhtrA* following entry into the mid-log phase. The mutant cells became elongated with sign of membrane blebs formation, an indication of cell death and lysis. Figure 6B showed a snapshot of cells harvested at stationary phase from 34°C and 37°C. Compared to the WT and *ΔhtrA^com^,* the mutant showed signs of distress with massive cell debris likely resulted from extensive cell lysis. This observation was exacerbated further at 37°C (**Fig. 6B**). Collectively, these results indicate that BbHtrA is critical for maintaining normal cell growth and morphogenesis especially at higher growth temperatures.

### The *ΔhtrA* mutant has attenuated motility and chemotaxis

Previous studies from our group and others identified the unpolymerized flagellin protein FlaB (40) and the chemotaxis phosphatase CheX (32) as the substrate for BbHtrA. In addition, over-expression of BbHtrA impaired *B. burgdorferi* motility, further indicating a role for BbHtrA in the motility and chemotaxis of *B. burgdorferi* (39). However, these data were obtained either using *in vitro* proteolytic analysis or from a BbHtrA over-expressed strain; thus, the exact role of BbHtrA on the spirochete motility and chemotaxis still remains elusive. To address this issue, we examined the swimming behavior of our mutant using swimming plate assay and bacterial cell tracking analysis as described previously (51). Swimming plate analysis showed that *ΔhtrA* is attenuated in swim ring formation on semisolid agar (14.5 ± 0.645 mm, n=4 plates) as compared to the WT (28.5 ± 1.19 mm, n=4 plates) and *ΔhtrA^com^*(27.5 ± 2.102 mm, n=4 plates). The swim ring formed by *ΔhtrA* was slightly larger than the non-motile *flaB* mutant (*ΔflaB*, 8.5 ± 0.289 mm), which serves as a reference for the size of the initial inoculum (**Fig. 7A**). This result reveals that *ΔhtrA* has attenuated motility as it failed to swim out from the initial inoculation site and formed swim rings that are only ∼50% in diameter of the WT and complemented strain. To further substantiate this result, we tracked WT, *ΔhtrA*, and *ΔhtrA^com^* cells and measured their cell velocities (**Fig. 7B**). The result showed that WT and *ΔhtrA^com^* cells are highly motile in 1% methylcellulose (videos 1 and 2) and reached 9.5 ± 2.53 µm/second (n=40 cells) and 10.91 ± 3.34 µm/second (n=34 cells), respectively. By contrast, *ΔhtrA* mutant cells were wiggling and unable to displace in 1% methylcellulose (**Fig. 7B** and video 3).

**Figure 7.**
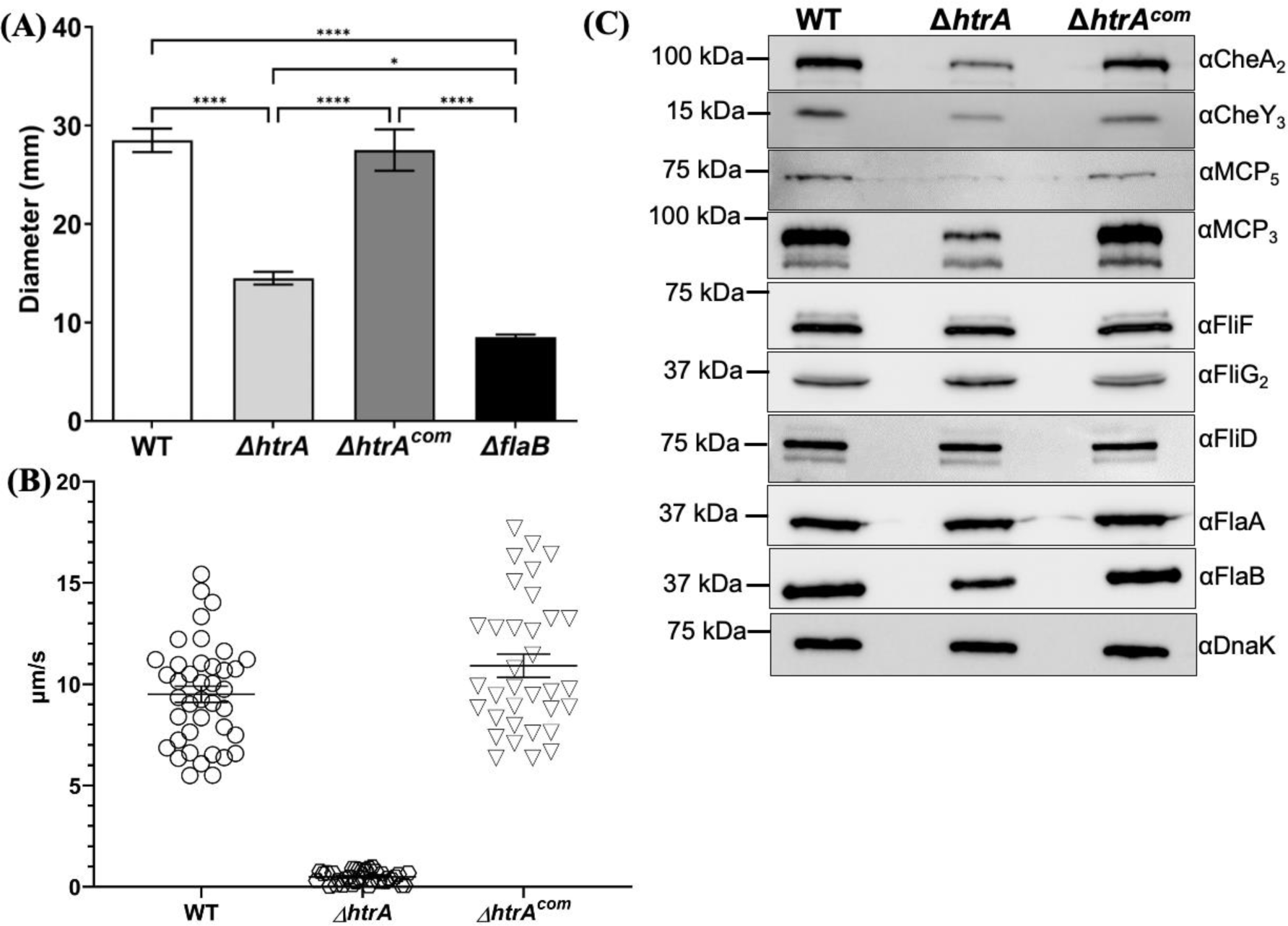
Deletion of BbHtrA represses spirochete motility and chemotaxis. **(A)** Swimming plate assays. The assay was carried out on 0.35% agarose containing 1:10-diluted BSK-II medium as described previously (53). The *flaB* mutant, *ΔflaB*, a previously constructed nonmotile mutant (52), was included to determine the sizes of inocula. The data are presented as mean diameters (in millimeters) of rings ± SEM for four plates. All assays were repeated at least twice with independent replicates and representative data are shown here. Statistical analysis was performed using ANOVA, followed by Tukey’s test for multiple comparisons, with a significance level of *P* value < 0.05. **(B)** Bacterial tracking analysis. The swimming velocities of WT, *ΔhtrA* and *ΔhtrA^com^*were measured using a computer-based motion tracking system as previously described (68). At least 30 cells were tracked and the swimming velocities were measured. **(C)** Immunoblotting analysis using different antibodies against select motility and chemotaxis-related proteins. Same amounts of WT, *ΔhtrA* and *ΔhtrA^com^*whole cell lysates were analyzed on SDS-PAGE and probed with antibodies against four flagellar proteins, including, FliF, FliG_2_, FlaA, and FlaB, and four chemotaxis proteins, including MCP_3_, MCP_5_, CheA_2_, and CheY_3_. DnaK was used as loading control.

### Deletion of BbHtrA affects the level of several flagellar and chemotaxis proteins

The periplasmic flagella (PFs) of *B. burgdorferi* have both the skeletal and motility functions (52). The altered motility and morphology phenotypes displayed by *ΔhtrA* suggest a possible linkage between BbHtrA and motility/chemotaxis. To test this possibility, we examined the expression level of several flagellar and chemotaxis proteins where antibodies are readily available in our laboratory using immunoblots. The expression of several flagellar proteins such as the MS-ring protein FliF, the motor switch protein FliG_2_, the flagellar filament cap protein FliD, and the flagellar sheath protein FlaA, were not affected in *ΔhtrA* (**Fig. 7C**). However, there was a slight decrease of FlaB level in *ΔhtrA* as compared to the WT and *ΔhtrA^com^*strain (**Fig. 7C**). Four chemotaxis proteins, including CheA_2_, CheY_3_, MCP_3_ and MCP_5_, were significantly lower in *ΔhtrA* and successfully restored in *ΔhtrA^com^*, which could have resulted in the attenuated chemotaxis of *ΔhtrA* as observed in the swimming plate assays given the essential role of CheA_2_ and CheY_3_ in the chemotaxis of *B. burgdorferi* (53–55).

### Deletion of BbHtrA disrupts the flagellar homeostasis

The above immunoblots revealed that deletion of *htrA* led to reduction in FlaB level but had no impact on other flagellar related proteins (**Fig. 7C**), which cannot explain why the *ΔhtrA* mutant fails to displace in 1% methylcellulose. To answer this question, cryo-ET was performed on *ΔhtrA* and *ΔhtrA^com^* strains. The result showed that roughly 60% of the *ΔhtrA* cells (20 out of 33 cells) contained shorter flagellar filaments, some of which are misoriented, pointing away from the cell center, and failed to form a ribbon-like structure (**Fig. 8A, B, & video 4**). Complementation with a functional BbHtrA successfully restored the normal flagella which form a tight ribbon wrapping around the cell body extending toward the cell center (**Fig. 8C, D**). Intriguingly, one of the mutant cells was found to contain an atypical flagellum encased within a 29.8 nm membranous sheath that travels from the periplasmic space into the cytoplasm (**Fig. 8E-H**). To the best of our knowledge, this abnormal flagellum has not been reported before in any *B. burgdorferi* mutants.

**Figure 8.**
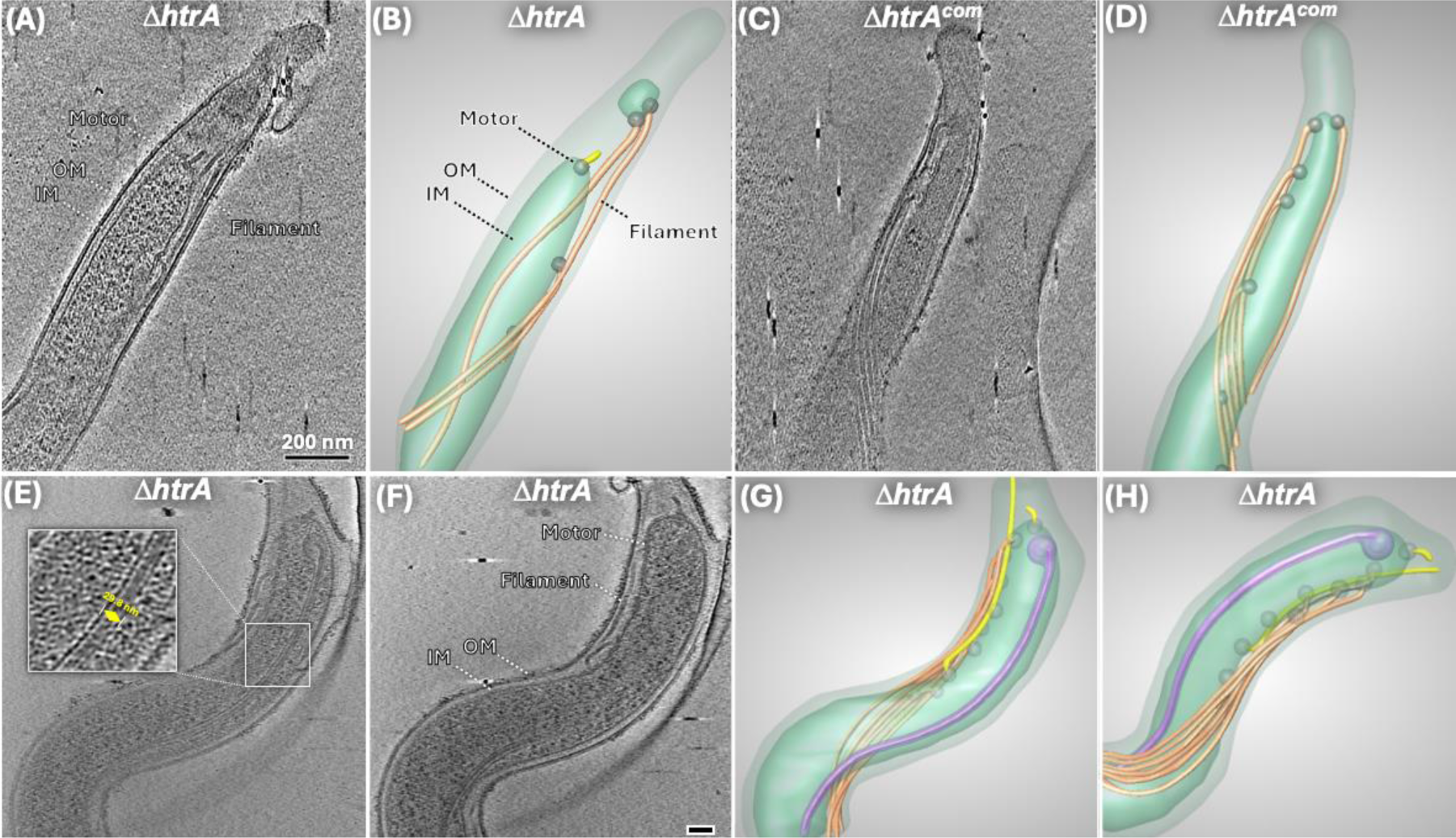
*ΔhtrA* forms short and misoriented flagellar filaments. 20 of 33 *ΔhtrA* cells examined contain defective flagella (short and misorientation). **(A** & **B)** are representative images showing the abnormal flagellum (yellow) and normal flagella (orange) in *ΔhtrA*. **(C** & **D)** Complementation of *ΔhtrA* restored the formation of normal flagella (orange). (**E** & **F**) Two different slices from one tomogram of a *ΔhtrA* mutant cell, respectively. An internal flagellum is encased within a membranous sheath with diameter of ∼29.8 nm. **(G** & **H)** Two different views of the 3D segmentation show the internal flagellum (purple) that traversed from the periplasmic space crossing into the inner membrane (IM).

### Deletion of BbHtrA leads to reduced BosR and RpoS

Mouse infection result showed that *ΔhtrA* was non-infectious and complementation using the native promoter successfully restored its infectivity (Table 1). This result supports the essential role of BbHtrA in the pathogenicity of *B. burgdorferi*, consistent with the observation made by Ye *et al.* (38). *B. burgdorferi* encodes several key transcriptional regulators such as BosR and RpoS (56–59) that govern the expression of surface proteins essential for host infectivity including OspC, BBK32, and DbpB/A (60–63). Immunoblots were performed to determine if deletion of BbHtrA leads to altered expression of regulatory proteins and subsequently antigen expression. Since *ΔhtrA* is significantly attenuated in growth at elevated temperature and reduced pH (i.e., 37°C/pH 6.8), we opted to culture the cells under laboratory condition (34°C/pH 7.4) to allow the mutant to achieve stationary growth phase. Immunoblotting analyses revealed that BosR expression was significantly reduced in the absence of BbHtrA as compared to the WT and the complemented strain (**Fig. 9A, B**). As expected, the levels of RpoS and RpoS-regulated proteins, such as BBK32, OspC, and DbpA, were also significantly downregulated in *ΔhtrA* as compared to the WT which were successfully restored in *ΔhtrA^com^*. The level of P66, which is not regulated via RpoS, remained relatively unchanged (64). Follow up qRT-PCR analysis revealed that the downregulation of BosR mainly occurred at the transcriptional level., e.g., *bosR* transcript was only expressed at ∼50% of the wild-type level which was restored in *ΔhtrA^com^* (**Fig. 9B**). Given that BosR positively regulates the transcription of RpoS (59) and the stability of *rpoS* transcript (65), in accordance with this, we saw a slight reduction in *rpoS* transcript in *ΔhtrA* but it was not statistically significant. However, the transcripts of RpoS-regulon, including *ospC* and *dbpA*, were significantly lower in *ΔhtrA* and were restored in *ΔhtrA^com^* (**Fig. 9B**), consistent with the immunoblot data (**Fig. 9A**).

**Figure 9.**
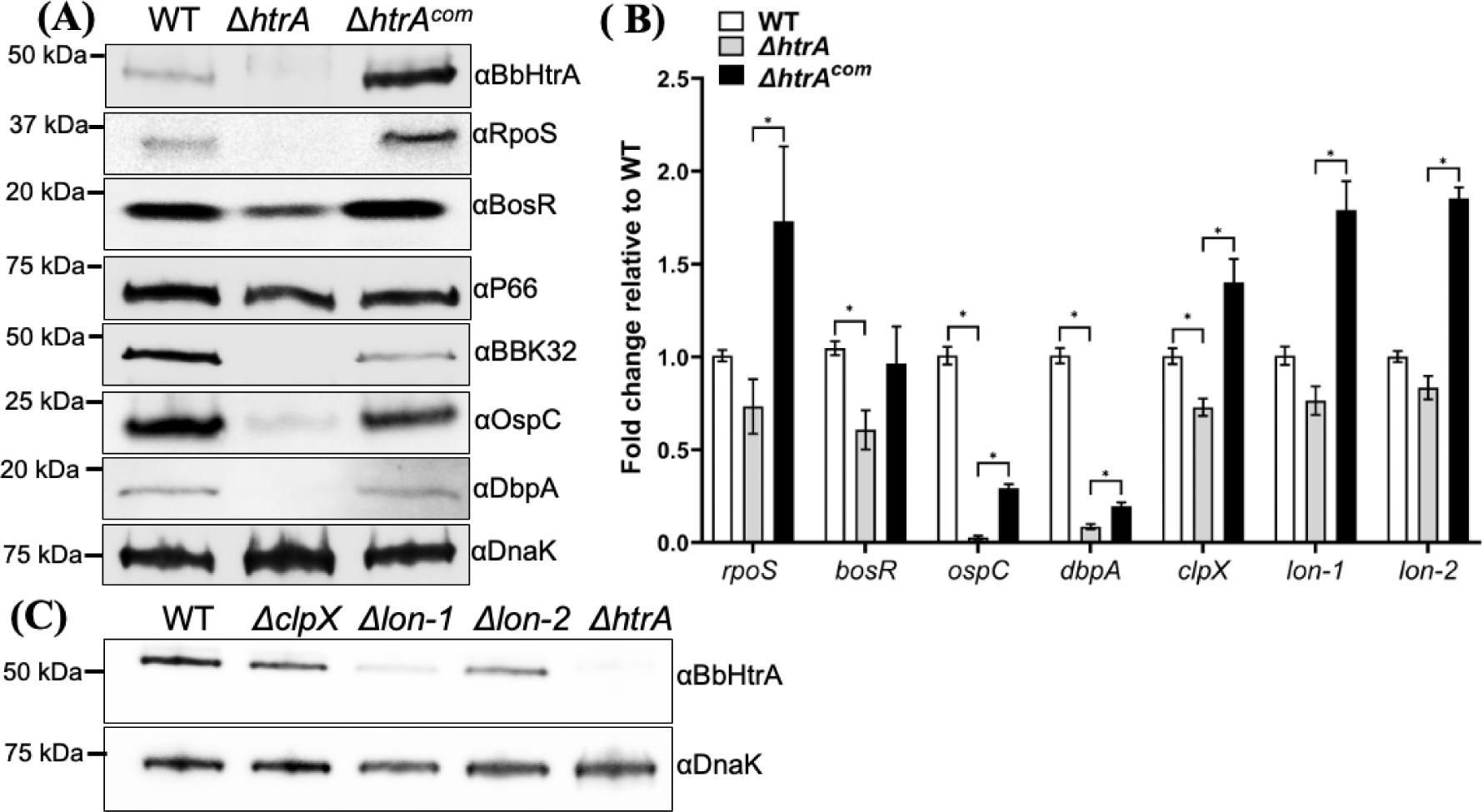
BbHtrA positively regulates BosR. **(A)** *B. burgdorferi* strains (WT, *ΔhtrA*, and *ΔhtrA^com^*) were cultivated at 34°C/pH 7.4 and harvested at stationary phase (∼10^8^ cells/ml) for immunoblotting analysis (A), and qRT-PCR analysis (B). **(A)** Immunoblotting analysis of WT, *ΔhtrA*, and *ΔhtrA^com^* samples using specific antibodies against BbHtrA, RpoS, BosR, P66 (98), BBK32 (99), OspC, and DbpA (73). DnaK was used as an internal control, as previously described (85). **(B)** Detection of *clpX*, *lon-1*, *lon-2*, *bosR*, *rpoS, ospC* and *dbpA* transcripts in WT, *ΔhtrA* and *ΔhtrA^com^* using qRT-PCR. The *dnaK* transcript was used as an internal control. Experiments were repeated at least three times with independent replicates. Data from five replicates are expressed as mean fold change relative to WT ± SEM. The significance of the differences between experimental groups was evaluated using multiple *t*-tests (*P* < 0.05), with correction for multiple comparisons performed using the False Discovery Rate (FDR) method. An asterisk (*) denotes a statistically significant discovery. **(C)** Detection of BbHtrA protein in *ΔclpX*, *Δlon-1*, and *Δlon-2* mutant. Equal amounts of whole cell lysates were analyzed on SDS-PAGE followed by immunoblotting analysis to detect BbHtrA level in the mutants harvested at mid-log phase. *ΔhtrA* was included as a control. DnaK was used as a loading control. Experiments were repeated at least twice with independent replicates.

### Deletion of BbHtrA leads to dysregulated protease expression

Recent studies showed that the expression of BosR and RpoS can be modulated by proteases, such as Lon proteases and ClpX (66–68). In addition, ClpXP protease is a known regulator of RpoS stability in other bacteria, such as *E. coli* (69–71). We found that the transcripts of *clpX* and two Lon proteases *lon-1*, and *lon-2* were significantly reduced in the absence of BbHtrA and were successfully restored in *ΔhtrA^com^* (**Fig. 9B**). This result implies that disruption of a single protease can have unexpected impact on the expression of other proteases in *B. burgdorferi*. Consistent with this proposition, immunoblotting analysis revealed that the level of BbHtrA in *ΔclpX*, *Δlon-1*, and *Δlon-2* mutants was altered to different extent in relative to that of the WT (**Fig. 9C**).

## DISCUSSION

Although the regulation of gene expression in *B. burgdorferi* has been studied extensively, majority of the known modulation occurs at the transcriptional and post-transcriptional level (14, 48, 65, 72–74). Proteases are a diverse group of enzymes that catalyzes the proteolytic degradation of select proteins within a cell to maintain protein homeostasis and cell viability (19). The genome of *B. burgdorferi* encodes for several proteases including the Lon protease family, ClpX, and the HtrA serine protease family (75). Recent studies on Lon proteases and ClpX suggest that proteolytic enzymes also contributes to the pathogenicity of *B. burgdorferi* by modulating the expression of select virulence factors and regulators (66–68). The function and activity of BbHtrA have been examined by several groups over the past decade (32–40). *B. burgdorferi* HtrA activity has been shown to be essential for the processing of several virulence factors including P66 (36) and BB0323 (38). However, the major shortcomings from the previous studies were the lack of a full phenotypic complementation and the use of an over-expression strain rather than a mutant to study the function of BbHtrA. The focus of our current study is to fill the gaps in the previous findings and attempts to decipher the regulation and contribution of BbHtrA to the pathophysiology of *B. burgdorferi*.

Complementation of *ΔhtrA* was as a major hurdle in studying the function of BbHtrA as attempts using *flaB* or *flgB* promoter-driven constructs failed to restore the infectivity of *ΔhtrA* in mice (Table 1) (38) even though the protein expression of BbHtrA was fully restored (**Fig. 1B**). This result indicates that the expression of BbHtrA requires a tight regulation *in vivo*. When we analyzed the genomic sequences around *htrA* (*bb0104*), we were unable to determine if *htrA* was part of a large gene operon or an orphan gene given the overlapping N-terminal sequences with *bb0105*. Using 5′-RACE analysis, we successfully mapped the TSS of *htrA* and discovered that the native promoter of *bb0104* appears to be located within the *orf* of *bb0105* (**Fig. 2**). However, we were unable to establish the class of this promoter based on the promoter elements as neither conserved -10 or -35 region can be identified using promoter prediction software (data not shown). In *ΔbosR* and *ΔrpoS* mutants, the promoter activities of *htrA* were significantly reduced in comparison to the WT but remained above background level across all time points. The common theme among the two mutants is the absence of the alternative sigma factor RpoS (57–59). Therefore, it is likely that BbHtrA expression is regulated by RpoS, either directly or indirectly, in response to certain stimuli or stress signals, given that HtrA enhances cell survival under stressful conditions in other bacteria species (76–80). As *B. burgdorferi* transition from the log- to stationary phase, we observed a gradual increase in the promoter activity of *htrA* which coincided with the increased in environmental stresses (e.g., high temperature, increase cell density, and reduced pH). Thus, the induction of BbHtrA expression is most likely needed to help the spirochetes survive under hardship conditions (**Fig. 3**). This is also in accordance with the mice infection study where RpoD (σ^70^) -driven constitutively active *flaB* and *flgB* promoter constructs failed to restore the infectivity of *ΔhtrA* (**Table 1**) because these promoters likely have different regulatory mechanism and expression pattern compared to the native promoter of *htrA*. Follow up immunoblotting analysis confirmed that several key virulence factors are differentially expressed in the four complemented strains, further emphasizing the significance of complementation using the native promoter (**Fig. 4**). Given the regulatory significance of the native promoter on BbHtrA expression during the life cycle of *B. burgdorferi*, in addition to BosR/RpoS, we reasoned that additional regulatory elements might be involved in the control of BbHtrA expression. Based on our luciferase reporter assay, the activity of *htrA* native promoter was only reduced but not abolished in the absence of BosR/RpoS (**Fig. 3**). Along with the lack of a conserved σ^70^ - 10 and -35 elements upstream of the TSS that was mapped (**Fig. 2A**), it remains a mystery how the promoter of *htrA* is activated and regulated. We performed promoter pulldown assays in order to identify potential regulatory element(s) associated with the native promoter region of *htrA* but was unsuccessful.

Using WT cultivated under various conditions, we discovered that the level of BbHtrA is differentially expressed (**Fig. 5**). Previous study indicated that the level of *htrA* transcript is highly elevated during stationary growth phase (38) which is consistent with the reporter assay result in WT where the strongest signal is detected when cells entered the stationary growth phase (**Fig. 3A**). Upon examining the protein level, we discovered that BbHtrA expression was significantly higher under tick-like environment when compared to the condition mimicking mammalian hosts. This observation suggests that BbHtrA may have a more significant role in the tick vector, perhaps functioning to suppress unnecessary anabolic processes so that the spirochetes can remain dormant while waiting for the next bloodmeal and transmission. Future study focusing on the role of BbHtrA in ticks is necessary in order to fully comprehend the significance of BbHtrA in the arthropod vector. In the study conducted by Ye *et al.*, it was reported that BbHtrA protein level was significantly increased when cultivated under host-like condition when compared to laboratory condition (38). This is in stark contrast to our observation where the expression of BbHtrA at 37°C was lower than 34°C (**Fig. 5**). This discrepancy could be due to the housekeeping gene used for the normalization analysis. Given that FlaB subunit is a substrate for BbHtrA (40) along with our observation that deletion of BbHtrA led to reduced FlaB level and flagellar assembly (**Figs. 7 & 8**), we reasoned that FlaB may not be the best housekeeping candidate for normalization. This could explain the discrepancy in relative expression of BbHtrA observed between Ye *et al.* and our study.

Given that HtrA is essential for cells to thrive under stressful conditions, we expect that the depletion of BbHtrA would compromise cell ability to survive under stress, such as stationary growth. By comparing the growth pattern of WT, *ΔhtrA,* and *ΔhtrA^com^* under various culture conditions mimicking the UF to mammalian host, we found that *ΔhtrA* has a significantly reduced growth rate as compared to the WT (**Fig. 6A**). The mutant failed to reach the same high cell density as the WT under all examined conditions with the most significant defect observed at elevated temperature mimicking the host environment, e.g., nearly 1 log difference in cell density was observed under this condition. Complementation of *ΔhtrA* fully restored the growth defect in *ΔhtrA* which strongly suggests that the growth defect is due to the absence of BbHtrA in the mutant (**Fig. 6A**). Microscopic examination of the mutant at stationary growth phase revealed that *ΔhtrA* cells have lost the regular spiral shape and are more elongated with membrane blebs observed on the cell surface as compared to the WT and *ΔhtrA^com^*. This alteration was exacerbated at 37°C as compared to 34°C growth condition (**Fig. 6B**). In addition, a high level of cell debris can be seen floating in the mutant culture, indicating an enhanced cell death and cell lysis event, which was corroborated with propidium iodide dead cells staining (**Fig. 6C**). This observation is consistent with the report by Ye *et al.* where deletion of BbHtrA in *B. burgdorferi* 297 strain also resulted in cell growth arrest and change in cell morphology at high temperature (38). Under a stressful growth condition, BbHtrA activity is likely necessary to assist with protein homeostasis within bacterial cells, such as to ensure proper protein folding and quality control. Without BbHtrA, toxic accumulation of misfolded protein can interfere with the regular cellular function leading to reduced fitness and subsequent cell death (21, 24).

Using a BbHtrA over-expressed strain, Coleman *et al.* showed that the ability of *B. burgdorferi* to form swim ring is significantly repressed, suggesting a potential connection between BbHtrA and spirochete motility/chemotaxis. We repeated the same experiment using our *ΔhtrA* and found that similar to the over-expression strain, depletion of BbHtrA resulted in the same phenotype where the mutant is attenuated in its ability to swim out in agar plates. The diameter of swim rings formed by *ΔhtrA* is larger than the non-motile Δ*flaB* but 50% smaller than that formed by both the WT and *ΔhtrA^com^*, indicating that the motility of *ΔhtrA* is significantly compromised in the absence of BbHtrA (**Fig. 7A**). Follow up immunoblotting analysis targeting various motility and chemotaxis-related proteins revealed that the expression of the MS-ring protein FliF, the motor switch protein FliG_2_ and the flagellar sheath protein FlaA remained unchanged (**Fig. 7C**), suggesting that the formation and assembly of flagellar motors is not affected in *ΔhtrA*, which was substantiated by the cryo-ET analysis (**Fig. 8**). Previous study suggests that FlaB is a substrate of BbHtrA. Therefore, we expect that deletion of *htrA* would increase the level of FlaB. However, an opposite effect was observed, e.g., the level of FlaB in the *ΔhtrA* mutant decreased by ∼50% (**Fig. 7C**). Our previous study shows that CsrA, a small RNA binding protein, inhibits FlaB translation (46). It is possible that deletion of BbHtrA increases the level of CsrA which in turn represses the expression of FlaB. We will investigate this possible mechanism in our future study. Two key chemotaxis proteins and two methyl-accepting chemotaxis proteins were significantly reduced in *ΔhtrA*. The low level of CheA_2_ and CheY_3_ (**Fig. 7C**), along with the formation of defective flagella (**Fig. 8**), could have contributed to the reduced swim ring formation in *ΔhtrA* as they are essential for the chemotaxis and motility of *B. burgdorferi* (53, 54). Additionally, impaired chemotaxis/motility could have contributed to the inability of *ΔhtrA* to establish infection in host, as chemotaxis and motility are essential for the infectious lifecycle of *B. burgdorferi* (81, 82).

In addition to motility and chemotaxis proteins, we have also examined several regulators and virulence factors essential for the survival and infectivity of *B. burgdorferi* in mice (**Fig. 9A**). We found that BbHtrA positively regulates BosR expression as deletion of BbHtrA led to reduced transcription of *bosR* and subsequently BosR protein expression while complementation restored the expression of BosR (**Fig. 9A, B**). Lower BosR in *ΔhtrA* impaired RpoS expression which in turn reduced the expression of RpoS-regulon genes such as OspC and DbpA. The data from qRT-PCR and immunoblot analyses (**Fig. 9A, B**) are consistent with the mouse infection study where dysregulated BosR-RpoS pathway resulted in the failure of *ΔhtrA* to establish an infection in the host (Table 1). BbHtrA is a protease and cannot directly control the transcription of *bosR*.

Therefore, it is possible that deletion of BbHtrA causes dysregulation of other regulators which in turn affects the transcription of *bosR*. Alternatively, deletion of BbHtrA may have disturbed the expression and activity of other proteases, such as ClpX, Lon-1, and Lon-2, which then causes systemic dysregulation of genes and proteins in *ΔhtrA* and vice versa (**Fig. 9C**). *B. burgdorferi* Lon proteases have been shown to affect BosR or RpoS-regulon (66, 67). Latest data from our laboratory also implicated a role for ClpX in the expression of RpoS (68). Consistently, our qRT-PCR data showed that the transcript of these three proteases is significantly reduced without BbHtrA and was successfully restored in *ΔhtrA^com^* (**Fig. 9C**). The reduced expression of other family of proteases could have further exacerbated the phenotype of *ΔhtrA*. Lastly, we also noted that in the various protease mutants available in our laboratory, the expression level of BbHtrA is affected, e.g., its level was found to be repressed in Lon proteases deficient mutants especially in *Δlon-1* (**Fig. 9C**).

In summary, our findings in this report aptly complement the current knowledge gap in understanding the contribution of BbHtrA in the pathogenicity of *B. burgdorferi*. BbHtrA contributes to the virulence of *B. burgdorferi* by impacting motility, chemotaxis, and the expression of several key regulatory and virulence factors in addition to the processing of BB0323 pre-protein. To fully appreciate the significance of this protease in the life cycle of the spirochete, future proteomic analysis examining the differentially expressed proteins between WT, *ΔhtrA,* and the fully complemented *ΔhtrA*^com^ strain is necessary to shed additional light in identifying novel target proteins that are affected by BbHtrA.

## MATERIAL AND METHODS

### Bacteria strains and culture conditions

A low-passage, virulent *B. burgdorferi* strain B31 A3-68 (83) was used in this study. *BosR* mutant (Δ*bosR*) (59), *rpoS* mutant (Δ*rpoS*) (84), *lon-1* mutant (Δ*lon-1*) (67), and *lon-2* mutant (Δ*lon-2*) (66) were kindly provided by Z. Ouyang (University of South Florida). *ClpX* (Δ*clpX*) mutant was previously constructed (68). *B. burgdorferi* wild-type (WT) and mutant strains were cultivated in Barbour-Stoenner-Kelly (BSK-II) medium as previously described (85) with appropriate antibiotics as needed: 300 μg/ml for kanamycin, 50 μg/ml for streptomycin, and/or 40 μg/ml for gentamicin. Cells were maintained at either 34°C/pH 7.4, or 37°C/pH6.8, in the presence of 3.4 to 5 % CO_2_. The antibiotic concentrations used for *E. coli* selection were as follows: kanamycin 50 μg/ml, spectinomycin 50 μg/ml, and ampicillin 100 μg/ml.

### Construction of *htrA* (*bb0104*) in-frame deletion mutant and its isogenic complemented strains

The deletion mutant of *bb0104* (*ΔhtrA*) was constructed in B31 A3-68 as previously described (40). To complement *ΔhtrA*, the intact *bb0104* gene was PCR amplified using primer pair P_3_/P_4_ along with *flaB* promoter (P_1_/P_2_), *flgB* promoter (P_5_/P_6_) or the native promoter of *bb0104* (P_7_/P_8_). The resulting promoter amplicons were PCR ligated to *bb0104* gene and then cloned into either pBSV2G (51) or pBBE22G (85) shuttle vectors at the BamHI and PstI sites, yielding four complementation constructs of pflgBbb0104/pBSV2G, pflaBbb0104/pBSV2G, pflgBbb0104/pBBE22G, and p104bb0104/pBSV2G (**Fig. 1A**). To complement *ΔhtrA*, these four constructs were transformed into the mutant cells, respectively. Complemented clones were selected on semi-solid agar plates containing both streptomycin and gentamicin. The primers for constructing the *ΔhtrA* complemented strains are listed in Table 2. PCR was used to detect the full plasmid profile of *ΔhtrA* and its isogenic complemented strains (**Table 1**) as previously described (86) to ensure retention of all necessary plasmids for mammalian infection study (**Fig. 1D**).

### Measuring the growth rates of *B. burgdorferi*

To measure the growth rates of the WT B31 A3-68, *ΔhtrA* and *ΔhtrA^com^* (an isogenic complemented strain of *ΔhtrA*), stationary-phase cultures were inoculated into fresh BSK-II medium to a final concentration of 1 x 10^5^/ml and incubated at 23°C/pH 7.4 (unfed tick condition, UF), 34°C/pH 7.4 (laboratory culture condition), or at 37°C/pH 6.8 (mammalian host condition). Bacterial cells in the cultures were enumerated every one to three days until reaching the stationary phase using a Petroff-Hausser counting chamber as previously described (87). Cells were counted with three biological replicates in three independent experiments; the results are expressed as the means ± standard errors of the means (SEM).

### Propidium iodide staining and cell imaging

To determine the percentage of dead cells, 10 µl of 600 µM propidium iodide was added to the cell culture prior to cell counting. The percentage of dead cells was enumerated using the ratio of propidium iodide positive cells over total cells. Images were taken using a Zeiss Axiostar plus microscope under 20 x objective lens and processed using Axiovision software (Zeiss, Germany) as previously described (88).

### Mouse infection studies

BALB/c mice at 6 to 8 weeks of age (Jackson Laboratory, Bar Harbor, ME) were used in the needle infection studies. All animal experimentation was conducted following the NIH guidelines for housing and care of laboratory animals and was performed in accordance with the Virginia Commonwealth University institutional regulations after review and approval by the Institutional Animal Care and Use Committees. The animal studies were carried out as previously described (81). Mice were given a single subcutaneous injection of 100 µL containing 10^5^ spirochetes and sacrificed at 3 weeks post infection. Tissues from the skin (inoculation site), ear, joint, heart, and spleen were harvested for re-isolation of spirochetes in BSK-II medium.

### 5′-Rapid Amplification of cDNA Ends (5′-RACE)

To determine the transcriptional start site (TSS) of *bb0104*, 5′ RACE was performed as previously described (50). In brief, wild-type B31 A3-68 cells were cultivated until late-log phase under laboratory culture condition. RNA was extracted using NucleoSpin RNA kit following the manufacturer’s instruction (Macherey-Nagel, Bethlehem, PA). 5′-RACE was carried out using SMARTer RACE 5’ Kit (Takara Bio USA, Mountain View, CA) following the manufacturer’s protocol. Primer used for the 5′-RACE is listed in Table 2.

### *β*-galactosidase assay

The upstream region of *htrA* was PCR amplified with primer pair P_8_/P_9_ (Table 2), generating an amplicon with engineered SmaI and BamHI cut sites at the 5’ and 3’ ends, respectively. The resultant amplicon was cloned into pGEM-T Easy vector (Promega, Madison, WI) and then released by SmaI and BamHI digestion. The released DNA fragment was cloned into pRS414 (40, 89), a *lacZ* reporter plasmid (a gift from R. Breaker, Yale University). As a positive control, *pflaB*, a previously constructed pRS414 vector containing the *flaB* promoter (90) was included (40). The resultant plasmids were transformed into the *E. coli* DH5α strain. Galactosidase activity was measured, as previously described (40). The results were expressed as the average Miller units of triplicate samples from three independent experiments ± standard errors of the means (SEM).

### Luciferase reporter assay

The native promoter of *htrA* was cloned into a luciferase construct, pJSB161 (49), at BglII and NdeI cut sites, using primer pair P_11_/P_12_. The obtained construct was transformed into B31 A3-68 (WT), *bosR* mutant (Δ*bosR*) (59), and *rpoS* mutant (Δ*rpoS*) (84), respectively. To monitor the activity of the *htrA* promoter, 10^6^ *B. burgdorferi* cells were inoculated into 5 mL of BSK-II medium and cultured under mammalian host conditions (37°C/pH 6.8). As a control, the WT carrying the empty vector pJSB161 was included. Bacterial cell densities of the cultures were measured every two days over twelve days using a Petroff-Hausser counting chamber. The luciferase activity was quantified as described previously (50). Briefly, 100 µL of bacterial cultures were inoculated into a 96-well plate. Fifty microliters of freshly prepared 2 mM D-luciferin substrate (Millipore-Sigma, Burlington, MA) was added to each well, mixed, and incubated for 1 minute, followed by reading using a Varioskan LUX multimode microplate reader (Thermo Fisher Scientific, Rockford, IL). Data were normalized to WT reading and expressed as normalized relative light units (RLU) per 10^5^ cells ± standard errors of the means (SEM).

### Bacterial motion tracking analysis and swimming plate assays

The swimming velocity of *B. burgdorferi* cells was measured using a computer-based motion tracking system as previously described (91). Briefly, log-phase *B. burgdorferi* cultures was first diluted (1:1) in BSK-II medium and 20 μl of the diluted cultures were mixed with an equal volume of 2% methylcellulose with a viscosity of 4000 cp (MC4000). Then *B. burgdorferi* cells were video captured with iMovie software on a Mac computer. Videos are exported as QuickTime movies and imported into OpenLab (Improvision Inc., Coventry, UK) where the frames were cropped, calibrated, and saved as LIFF files. The software package Volocity (Improvision Inc.) was used to track individual moving cells and cell velocities were calculated. For each bacterial strain, at least 20 cells were recorded for up to 30 sec. Swimming plate assays was performed using 0.35% agarose with BSK-II medium diluted 1:10 with Dulbecco’s phosphate-buffered saline (DPBS, pH 7.5), as previously described (52, 53, 92). WT, *ΔhtrA, ΔhtrA^com^*, and *flaB* mutant (*ΔflaB*), a previously constructed non-motile mutant (52) was included as a negative control to determine the initial inoculum size. The plates were incubated for 4–5 days at 34°C in the presence of 3.4% CO_2_. Diameters of the swim rings were measured and recorded in millimeters (mm). The average diameters of each strain were calculated from four independent plates. Statistical analysis was performed using One-way ANOVA, followed by Tukey’s test for multiple comparisons, with a significance level of *P* value < 0.05.

### RNA preparation and quantitative reverse transcription-PCR (qRT-PCR)

RNA isolation was performed as previously described (87). Briefly, *B. burgdorferi* strains were cultivated at 37°C/pH 6.8, and 10 ml of mid-log (∼10^6^ cells/ml) or stationary phase cultures (∼10^8^ cells/ml) were harvested for RNA preparation. Total RNA was extracted using TRIzol reagent (Thermo Scientific), following the manufacturer’s instruction. The RNA samples were then treated with DNase (Takara Bio USA) at 37°C for 2 hours to eliminate genomic DNA contamination. The resultant RNA samples were re-extracted using acid phenol-chloroform, precipitated in isopropanol, and washed with 70% ethanol. The RNA pellets were re-suspended in RNase-free water. cDNA was generated from total RNA using SuperScript IV VILO master mix (Thermo Fisher Scientific). qRT-PCR was performed using Fast SYBR™ Green Master Mix (Applied Biosystems, Foster City, CA) on a QuantStudio 3 real-time PCR system (Thermo Fisher Scientific). The chaperon gene (*dnaK*, *bb0518*), a housekeeping gene, was included as an internal control to normalize the qRT-PCR data. The results were expressed as the fold change relative to the WT. The primers used for qRT-PCR are either listed in Table 2 or described in our recent report (68). The significance of the differences between experimental groups was evaluated using multiple unpaired *t*-tests (*P* < 0.05) with correction for multiple comparisons performed using the False Discovery Rate (FDR) method. An asterisk (*) denotes a statistically significant difference.

### Gel electrophoresis and immunoblotting analysis

Sodium dodecyl sulfate-polyacrylamide gel electrophoresis (SDS-PAGE) and immunoblotting with an enhanced chemiluminescence detection method were carried out as previously described (87). Approximately 10-20 μg of bacterial whole cell lysates was loaded into each lane of 12% SDS-PAGE gels and transferred to polyvinylidene difluoride (PVDF) membranes (Bio-Rad Laboratories, Hercules, CA). The immunoblots were probed with antibodies against *B. burgdorferi* proteins of interest. Monoclonal and polyclonal antibodies were generously provided by the following investigators: monoclonal anti-FlaB (H9724) by A. G. Barbour (University of California, Irvine, CA), monoclonal anti-FlaA by B. Johnson (Center for Disease Control and Prevention, Atlanta, GA), monoclonal anti-DnaK by J. Benach (SUNY at Stony Brook, NY). Antibodies against BosR, P66, DbpA, and OspC are as described (68), polyclonal anti-RpoS was raised in-house, antibodies against *B. burgdorferi* HtrA, FliF, FliG_2_, MCP_3_, MCP_5_, CheY_3_ and CheA_2_ were described in our previous publications (40, 53–55, 88, 93). The immunoblots were developed using horseradish peroxidase secondary antibody with an ECL luminol assay. Signals were imaged using the ChemiDoc MP imaging system and quantified with Image Lab software (Bio-Rad Laboratories). At least three biological replicates of cells were harvested and used for immunoblot analysis.

To determine the expression of BbHtrA under different growth conditions, 10^5^ cells/ml of wild-type *B. burgdorferi* were inoculated into 10 ml of BSK-II medium (pH 7.4 for 23°C and 34°C, and pH 6.8 for 37°C). Cells were cultivated for 7–10 days and harvested at mid-log phase (∼10^6^ cells/ml) or at stationary phase (∼10^8^ cells/ml) and used for immunoblots. The immunoblot signals were quantified using Image Lab. Experiments were repeated twice and data is expressed as mean BbHtrA expression level relative to DnaK ± SEM.

### Cryo-ET sample preparation

*ΔhtrA* and *ΔhtrA^com^* samples were centrifuged at 1,500 × *g* for 10 minutes and resuspended in phosphate buffer saline (PBS). 10 nm gold tracer (AURION Immuno Gold Reagents & Accessories, The Netherlands) was used on bacterial samples at a ratio of 1:1 (V/V). Leica EM GP2 plunger (Leica Microsystems Inc., Deerfield, IL) was used for sample cryo-fixation. GP2 environmental chambers were set to 25°C and 95% humidity, and 5 μL of each *B. burgdorferi* samples were applied to the carbon side of the freshly discharged grid. After 30 seconds, the grids were blotted for 6 seconds from the backside and immediately plunged frozen in liquid ethane.

### Cryo-ET data collection

Frozen hydrated *B. burgdorferi* specimens were examined on a Glacios 200kV Transmission Electron Microscope (ThermoFisher) equipped with a K3 direct detection camera (Gatan Inc., Pleasanton, CA). SerialEM software and FastTomo script were used for tilt series data collection at low-dose mode, 13,500 × magnification, physical pixel size 3.25Å with 6 μm defocus. 33 images were recorded from -48° to +48° with 3° tilt increment. The total electron doses were ∼ 66 e^-^/Å^2^.

### Cryo-ET data analyses

Image drifting induced by electron beam was corrected by MotionCor2 (94). IMOD software (95, 96) was used to create image stacks and aligned all images in each tilt series by tracking with fiducial beads. 8 × binned tomograms were reconstructed using Tomo3D (97). In total, 110 tomograms were generated from *ΔhtrA* and 18 from *ΔhtrA^com^* strain, respectively. IMOD (95, 96) was then used to take snapshots from the tomograms. Dragonfly software (Vesion 2022.2, Comet Technologies Canada Inc., Quebec, Canada) was used to manually segment the features including the outer membrane, inner membrane, and flagella.

### Statistical analysis

For the swimming plates and motion tracking analysis, the results are expressed as means ± standard error of the mean (SEM). The significance of the difference between different experimental groups was evaluated with ANOVA (*P* value < 0.05).

## ACKNOWLEDGMENT

We thank Dr. Zhiming Ouyang for providing the *lon-1*, *lon-2*, *rpoS*, and *bosR* mutants, Dr. Melissa Caimano for providing OspC and DbpA antibodies, Dr. Jennifer Coburn for providing P66 antibody, and Dr. Janakiram Seshu for providing BBK32 antibody. This work was supported by funding from the National Institutes of Allergy and Infectious Diseases (AI078958 to C. Li; AI087946 to J. Liu, AI148844 to B. Crane and C. Li), National Institutes of Health (NIH).

